# Creation of MLPA: A Multi-level Digital Twin Framework for Personalized Cancer Simulation and Treatment Optimization

**DOI:** 10.1101/2024.09.13.612988

**Authors:** James Gu, Jake Y Chen

## Abstract

This research proposes a new personalized cancer modeling and treatment strategy: the Multi-level Parameterized Automata (MLPA), an innovative digital twin framework. The MLPA framework fully integrates macroscopic electronic health records (EHR) and microscopic genomic data for the first time, employing stochastic cellular automata to model tumor progression and treatment efficacy dynamically.

The multi-scale strategy effectively enables MLPA to simulate complex cancer behaviors, including metastasis and pharmacological responses, with remarkable precision. The validation using bioluminescent imaging from mice demonstrates MLPA’s exceptional predictive power, achieving a state-of-the-art R² value of 0.93, improvement over previously reported conventional models, and fitting within 95% confidence intervals.

The proposed MLPA framework accurately captures tumors’ characteristic S-shaped growth curve and shows high fidelity in simulating various scenarios, from natural progression to aggressive growth and drug treatment responses. MLPA’s ability to simulate drug effects through gene pathway perturbation, with predicted tumor cell counts being within a 2.5% equivalence margin interval, underscores its potential as a powerful method for precision oncology.

The proposed MLPA framework not only offers a more reliable platform for exploring personalized treatment strategies, but also potentially transforms patient outcomes by optimizing therapy based on individual human biological profiles. As clinically adopted technology progresses towards precision medicine, MLPA stands at the forefront, offering new possibilities in cancer simulation and treatment optimization. The code and imaging dataset used is available at https://github.com/alphamind-club/MLPA.

## Introduction

Based on the Center for Disease Control and Prevention’s recent report, cancer is the second leading cause of death in the United States, with 608,371 deaths recorded annually ^1^. By 2040, there will be a predicted 26.0 million cases of cancer in the United States alone, a massive increase from the 18.1 million diagnoses in 2022. Cancer is particularly dangerous due to its ability to evade the body’s immune system, while spreading to other organs through a process known as metastasis ^2^. Its rapid and uncontrolled cell growth can overwhelm normal tissues, disrupting vital functions and making treatment difficult, especially in advanced stages while also being difficult to detect in preliminary stages. This staggering growth of cancer cases highlights the urgent need for the development of more effective cancer prediction and treatment methods in today’s clinical workflows. Despite significant advancements in oncology, cancer’s inherent complexity and heterogeneity, marked by distinct genetic mutation patterns and diverse clinical presentations ^3–4^, often render traditional drug treatment strategies ineffective, leading to suboptimal patient outcomes ^5–6^. The challenge lies in developing personalized treatment approaches that account for the unique characteristics of each patient’s cancer ^7–8^.

In recent years, advancements in cancer prediction—aimed at simulating disease progression and optimizing treatment strategies—have primarily been driven by two key approaches: statistical modeling and machine learning techniques. The work of Mascheroni et al. stands out as an early effort to address data sparsity in personalized tumor growth prediction ^9^. By combining Bayesian and mechanistic modeling, they laid the groundwork for more robust cancer modeling, but their method still struggled with feature selection—a recurring challenge even in recent models. Lai et al. took a different route by using cellular automaton techniques for breast cancer therapy ^10^.

Their model is noteworthy for its scalability, but like many statistical approaches, it lacks generalizability across different tumor types, restricting its broader clinical utility. More recent research, such as Valentim et al., have continued to explore cellular automaton models. Although Valentim’s work offers an innovative way to simulate various tumor growth scenarios, the absence of real-world data calibration still raises concerns about accuracy when translating these simulations into clinical applications ^11^. Velasquez et al. compares parametric models for predicting overall survival in RET-altered solid tumors, showing superior accuracy for mathematical models but highlighting the need for further validation across tumor types ^12^.

Additionally, Wang et al. developed a prostate cancer prediction model using parametric statistical methods, though it lacks lifestyle and comorbidity data, limiting its overall utility ^13^. Despite these contributions, statistical methods often struggle to integrate diverse and large-scale datasets, which limits their predictive power.

On the other hand, it has been proven that the machine learning approach has an advantage over this limitation by having the capability to process large, complex datasets in tandem with multimodal data. Rabiei et al. and Tătaru et al. used Random Forest and Gradient Boosting Trees for breast and prostate cancer prediction, respectively, achieving around 80% accuracy with high sensitivity and specificity ^14–15^. However, their models’ lack of genomic or lifestyle data limits their generalizability, a recurring problem in this domain. Groheux et al. also explored machine learning in breast cancer prediction, yet similarly acknowledged that omitting genomic inputs constrained their model’s predictive power ^16^. Interestingly, Steyaert et al. pushed the boundaries of machine learning by incorporating both gene expression and histopathology data in their brain tumor prognosis models ^17^. This approach significantly enhanced survival predictions, offering a glimpse into the future of integrating multiple data types to improve accuracy. In contrast, Vale-Silva et al. integrated genomic data for long-term cancer survival prediction, but still faced performance variability across cancer types, highlighting that even models with comprehensive datasets need further refinement ^18^.

Moreover, very recent developments in digital twin technology, particularly from the Swedish Digital Twin Consortium, have sparked considerable interest in their potential to revolutionize personalized treatment strategies ^19^. Their work emphasized patient-specific models that simulate disease progression and treatment response, a marked shift from traditional predictive models.

Yet, the challenge remains in scaling this technology for broader healthcare applications. Benson et al. have taken this a step further by developing high-resolution patient models capable of computationally evaluating drug efficacy on an individual level, although achieving seamless integration of such models into clinical practice remains an ongoing challenge ^20^. Machado and Berssaneti’s review of digital twin applications reiterates the transformative potential in healthcare, though realizing fully functional, whole-body digital twins is still not yet achieved ^21^.

Although some progress has been achieved as mentioned above, there were still some research gaps that were identified through literature review. These research gaps are mainly highlighted as follows: 1) Previous approaches in cancer prediction focus on detection rather than modeling disease progression. While early detection is important, it is equally critical to forecast how cancer will evolve, as this informs treatment strategies and improves patient outcomes; 2) Despite achieving 80-90% accuracy levels, previously reported AI models function as black-boxes. They focus on performance metrics without providing transparency into the mechanisms driving cancer progression, limiting their clinical utility. Understanding key factors such as tumor growth, metastasis, and drug response are crucial for developing personalized and effective treatment plans; and 3) Currently reported studies heavily rely on a narrow set of data inputs, often missing crucial genomic and spatial transcriptomic data. This restricts their predictive power and limits their ability to offer personalized treatment strategies. Integrating multiple data types is desirable to further improve both accuracy and the relevance of cancer progression models, which would provide a more comprehensive and reliable view of the disease and its behavior over time.

To address the research gaps, this research focuses on the following areas: 1) Propose a novel computational MLPA framework. The proposed MLPA framework will fully integrate macroscopic EHRs with microscopic genomic data, which is now more accessible, to create a detailed, mechanism-based dynamic model of both the tumor microenvironment and the patient’s overall health profile; 2) Develop a strategic method utilizing stochastic cellular automata and multiple data modalities to simulate timeseries cancer imaging data. While stochastic cellular automata have been previously used to model tumor growth, the MLPA framework uniquely combines them with genomic, EHR, and imaging data to offer a more comprehensive and personalized view of cancer dynamics; and 3) Utilize Monte Carlo simulations to explore and optimize cancer prediction model parameters to reduce simulation error, instead of relying on traditional AI models which have black-box issues. By comprehensively incorporating a broader range of data types than existing models, the MLPA framework aims to improve predictive accuracy and provide a robust platform for optimizing individualized cancer treatment plans.

In summary, the proposed MLPA digital twin framework serves as a virtual testbed for simulating disease progression and evaluating therapeutic responses under various treatment regimens, providing an in-depth understanding of treatment outcomes, potential resistance mechanisms, and the optimization of personalized treatment plans. This research proposes, for the first time, the theoretical foundation of MLPA, its initial prototype, and its validation using a bioluminescent imaging mice dataset. The simulation results demonstrate MLPA’s ability to accurately predict tumor dynamics and evaluate personalized treatment strategies. By constructing this proof-of-concept cancer digital twin, this research aims to advance the frontier of precision medicine, enhancing the specificity and efficacy of oncological therapies.

The paper is organized as follows: The “Results” section presents a thorough analysis of the MLPA framework’s fitting, validation, and drug effect simulation, utilizing a bioluminescent imaging dataset of mice injected with MLL-rearranged leukemia and treated with DOT1L histones. Following this, the “Discussion” section delves into the implications and interpretations of these findings. The “Limitations” section openly addresses potential limitations and areas for enhancement. Finally, the “STAR METHODS” section outlines the proposed methodology and provides details on data and code availability.

## Results

The results presented here highlight MLPA’s versatility and predictive power in simulating tumor growth under various conditions, predicting tumor progression with high accuracy, and validating the effects of gene-targeted therapies. This section is organized into four subsections: 1) simulation of tumor growth in different scenarios; 2) predictive accuracy of MLPA compared to mice data; 3) validation of drug treatment effects through gene pathway perturbation; and 4) assessment of MLPA’s reliability in predicting time points beyond the training dataset.

### 1. Simulating Tumor Growth Across Control, Aggressive, and Treatment Scenarios

Figure 1 shows MLPA’s capability to simulate various scenarios of tumor progression, providing insights into various aspects of cancer growth and treatment responses. Importantly, it can simulate tumor dynamics across multiple time points, allowing for a more comprehensive understanding of disease progression and the evaluation of treatment efficacy over time.

**Figure 1:**
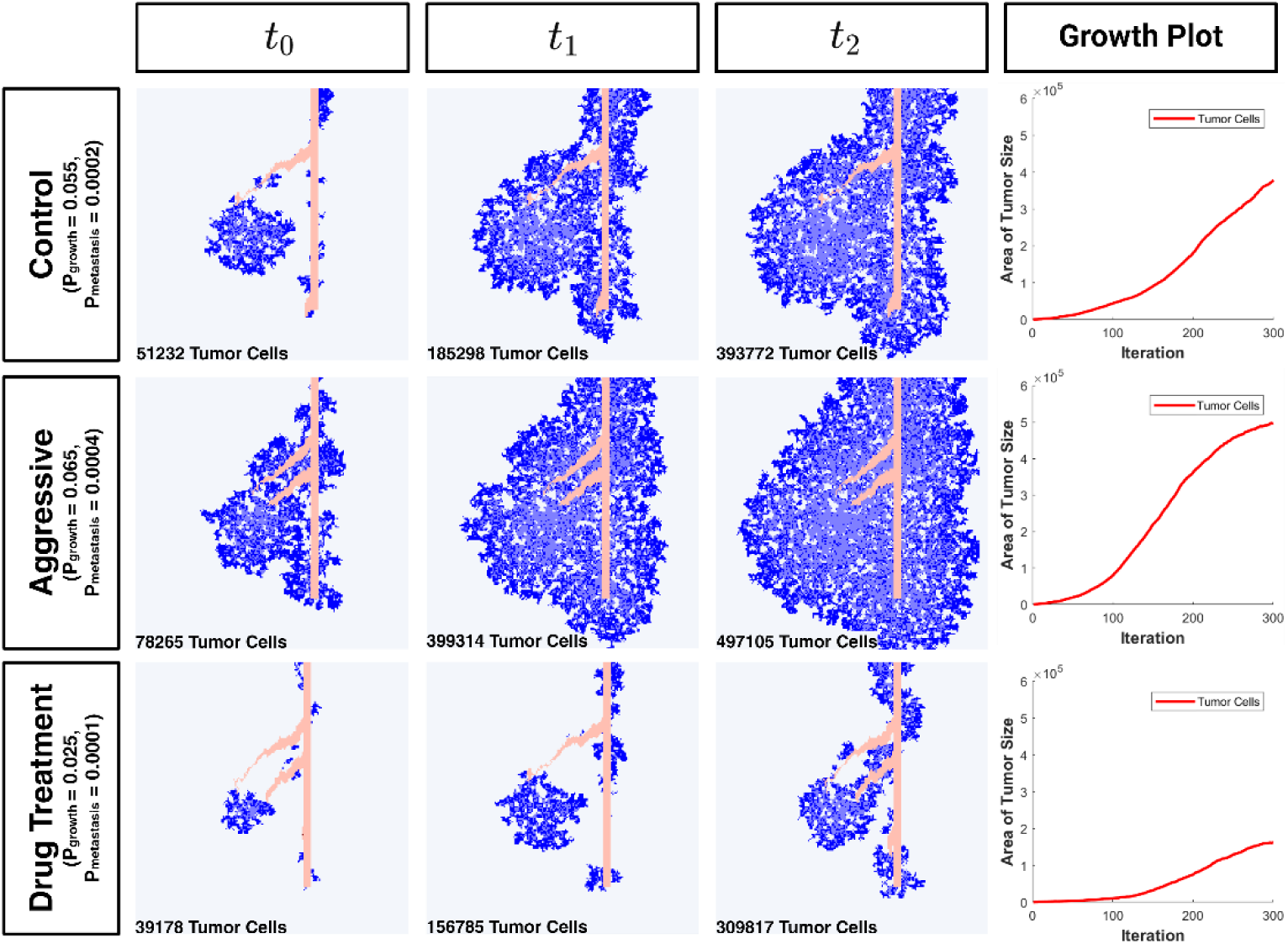
MLPA simulation of tumor growth under Control, Aggressive, and Drug Treatment conditions from t_0_ to t_2_. The Aggressive scenario shows increased growth due to upregulated cancer-promoting gene pathways, while the Drug Treatment scenario shows decreased growth due to downregulation. Iterations are defined as each individual passes of the code and tumor size is calculated by adding all the number of tumor cells grown.

In the control scenario (P_growth_ = 5.5 × 10^−2^ and P_metastasis_ = 2.0 × 10^−4^), the model simulated the natural progression of a tumor without intervention. Initially at t₀, the tumor was localized with limited spread. Over time, the tumor expanded steadily, showing increased density and area coverage. By t_1_ and t_2_, the tumor had grown significantly, occupying a larger area of the simulated tissue. This scenario validated the model’s ability to replicate natural tumor growth patterns in untreated conditions accurately.

The aggressive scenario (P_growth_ = 6.5 × 10^−2^ and P_metastasis_ = 4.0 × 10^−4^) simulated conditions where factors enhanced tumor growth and metastasis, with gene pathways in the PAGER3 database contributing to cancer advancement activated. By t_1_ and t_2_, there was rapid and extensive tumor growth, with significantly increased density and area coverage compared to the control scenario. The tumor size at t_2_in the aggressive scenario was approximately 1.26 times larger than in the control scenario, demonstrating the model’s ability to capture accelerated growth patterns.

The drug treatment scenario (P_growth_ = 2.5 × 10^−2^ and P_metastasis_ = 1.0 × 10^−4^) evaluated the efficacy of therapeutic interventions in controlling tumor growth, with gene pathways in the PAGER3 database contributing to cancer advancement inhibited. As time progressed to t_1_and t_2_, the tumor growth was noticeably slower, and the density was reduced compared to both the Control and Aggressive scenarios. At t_2_, the tumor size in the drug treatment scenario was approximately 0.38 times the size of the control scenario, illustrating the model’s capacity to simulate the effects of therapeutic interventions.

### 2. Validating Predictive Accuracy Using Sigmoidal Approximations

Building on the previous simulations, we evaluated the accuracy of MLPA’s predictions against real-world data from bioluminescent imaging in mice, specifically on the four testing nontreated mice subjects that were not used during the training phase. The MLPA framework was run four times, with the tumor pixel and high-density tumor pixel counts averaged. It was found that the results were able to be represented by a sigmoid function, with 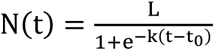, where N(t) is the number of cancer cells at time t, L represents the carrying capacity of tumor cells, k controls the growth of the cancer, and t_0_ is the time of the inflection point of the growth. The R^2^ value obtained was 0.93, achieved by comparing the MLPA predicted outputs with the curves fit to the bioluminescent mice imaging dataset. Other studies only achieved values of 0.80-0.90 ^14, 15^.

The sigmoid function is ideal for modeling tumor growth because it captures the initial slow growth phase, rapid exponential growth, and eventual plateau as the tumor reaches its maximum size, reflecting real-life conditions. The error bars in Figure 2A are the range of area values obtained from the testing split of the mice imaging data. Both the tumor pixel and high-density tumor predictions from MLPA fell within an acceptable range of error. MLPA predicted that the total carrying capacity would be at 1,006,630 cells, with the inflection point of growth being between 1 and 2 days. Notably, the model predicts a deceleration in cancer cell proliferation beyond Day 5, as indicated by the approaching plateau in the graph. This deceleration can be attributed to the tumor reaching spatial constraints within the host environment.

**Figure 2:**
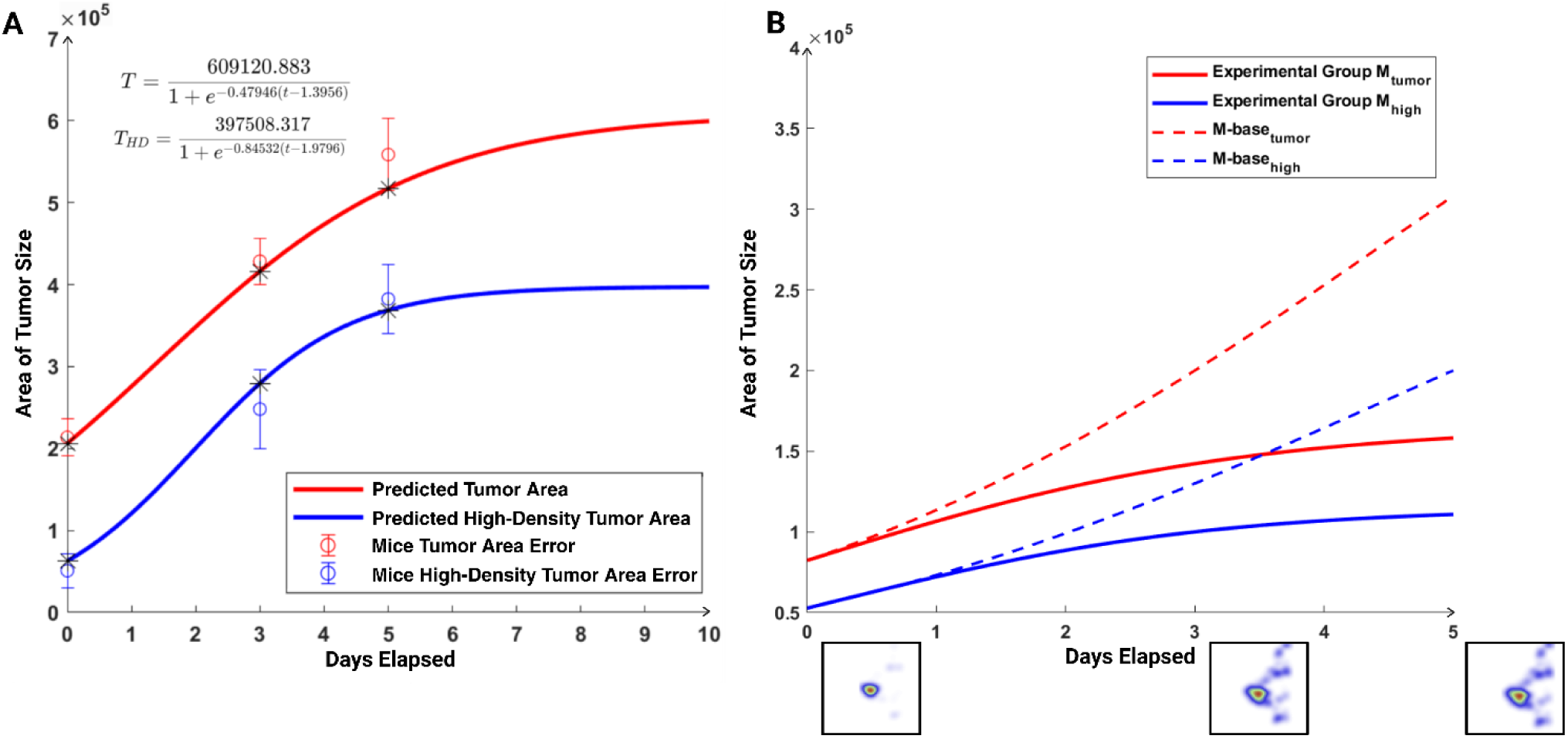
Sigmoid Fit and Experimental Group Predictions: A) Sigmoid fit comparing MLPA predicted tumor area (red) and high-density tumor area (blue) against observed tumor areas from mice data (open circles with error bars), with equations provided for both fits; and B) MLPA model predictions for the experimental group (4 treated mice subjects) treated with DOT1L inhibitor over five days. Solid lines represent predicted tumor areas with parameter perturbation from gene pathways enabled, while dotted lines show predictions without said perturbations.

While other models such as the Gompertz or logistic functions also capture nonlinear growth, the sigmoid model offers a balance between simplicity and precision, making it ideal for datasets with limited time points. For example, the Gompertz model often requires finer temporal granularity to estimate its asymmetric parameters reliably, while more complex models can risk overfitting or instability when applied to datasets with sparse observations.

### 3. Gene Pathway Targeting for Drug Efficacy Simulation in MLPA

To validate the framework’s treatment prediction using gene pathway perturbation, the MLPA framework was compared with the four mice treated with a small-molecule inhibitor of DOT1L, listed as the experimental group. The efficacy of this treatment was assessed by its ability to revert gene perturbations caused by the disease. After fitting the default parameters of the MLPA model to the experimental group (P_growth_ = 3.2 × 10^−2^ and P_growth_ = 2.8 × 10^−4^), specific gene pathways known to be inhibited by DOT1L were selected, which were then used to perturb the default model parameters as shown in Figure 2B. The following pathways from the CSV gene pathway file which would be inhibited during treatment were chosen as follows: 1) Nonsense Mediated Decay; 2) Chaperon Mediated Autophagy; 3) Regulation of TP53 Expression and Degradation; 4) TP53 Activity through Methylation; and 5) Aggrephagy. These pathways were inhibited so the result would counteract cancer progression, such as degrading potentially oncogenic proteins, maintaining protein homeostasis, enhancing tumor suppressor functions, and reducing proteotoxic stress.

A 95% confidence interval was created for the Day 5 experimental group predicted tumor pixel counts, and was compared to the Day 5 pixel counts of the drug treated mice in the dataset. The 95% confidence interval from the mice dataset for tumor pixel count was determined to be between 153,336 and 170,231 pixels when the predicted tumor pixel count was 158,125. The 95% confidence interval from the mice dataset for high-density tumor pixel count was determined to be between 97,215 and 120,958 pixels when the predicted high-density tumor pixel count was 110,807. Since the confidence intervals encapsulate the predicted values, they indicate a precise prediction.

To further validate these results, equivalence tests were conducted using the Two One-Sided Test procedure. For tumor pixels, the equivalence margin was set at ±3,953 pixels (2.5% of the observed value), with a 95% simulated predictive interval of [143,124, 167,784]. The difference between MLPA’s and the imaging dataset means was 2,670 pixels. For high-density tumor pixels, the equivalence margin was ±2,770 pixels (2.5% of the observed value), with a 95% simulated predictive interval of [101,545, 118,347]. The difference between MLPA’s and the imaging dataset means was 860 pixels. In both cases, MLPA’s predicted pixel counts fell within the intervals from the actual imaging dataset and with differences well within the defined equivalence margins. These findings indicate that the drug treatment significantly reduced tumor size and density, aligning closely with the MLPA framework’s predictions. The strong agreement within the confidence intervals underscores the model’s ability to accurately capture the effects of activated gene pathways, thereby validating the proposed approach to simulating drug effects through gene pathway perturbation. It was validated through experimental data that this alignment not only reinforces the model’s predictive accuracy, but also supports its potential utility in personalized medicine applications where understanding the impact of gene-targeted therapies is crucial.

In addition, when there is no gene pathway perturbation, as depicted in Figure 2B, we can clearly observe a significant deviation from the actual results. For example, the tumor pixel count predicted without pathway perturbation exceeded the observed value by 158,432 pixels (105% error) for Day 5, and the high-density tumor pixel count overestimated the observed value by 89,193 pixels (180% error). This discrepancy highlights the critical role of incorporating gene pathway perturbations in achieving accurate predictions. Without these perturbations, the MLPA framework does not capture the biological effects of the treatment, leading to vast overestimations of tumor progression and density.

### 4. Modeling Tumor Progression Through Time Point Prediction

Beyond validating drug effects, we also tested the framework’s ability to predict tumor progression across time points, even when some data points were withheld. Due to the specificity of the dataset required for MLPA, the mice dataset was limited to only three data points, which made it insufficient for this purpose, as hiding one point would leave too few for meaningful predictions. To overcome this limitation, an initial testing MLPA model was used with the initial parameters of P_growth_ = 4.5 × 10^−2^and P_metastasis_ = 3.0 × 10^−4^ to generate simulated imaging data for days 0, 2, 3, and 5. Then, the Day 2 cancer image was intentionally hidden, and a separate MLPA model was used to see if it could accurately generate the missing image without having access to the initial model’s parameters, only using the other three days’ imaging data to develop its own parameters, shown in Figure 3A. By effectively predicting the missing data for Day 2 on another model instance without knowing the initial parameters, MLPA shows its capability to generate continuous, reliable data, even when only a few time points are available. The second model adopted the parameters to P_growth_ = 4.45 × 10^−2^ and P_metastasis_ = 3.06 × 10^−4^ during the fitting process.

**Figure 3:**
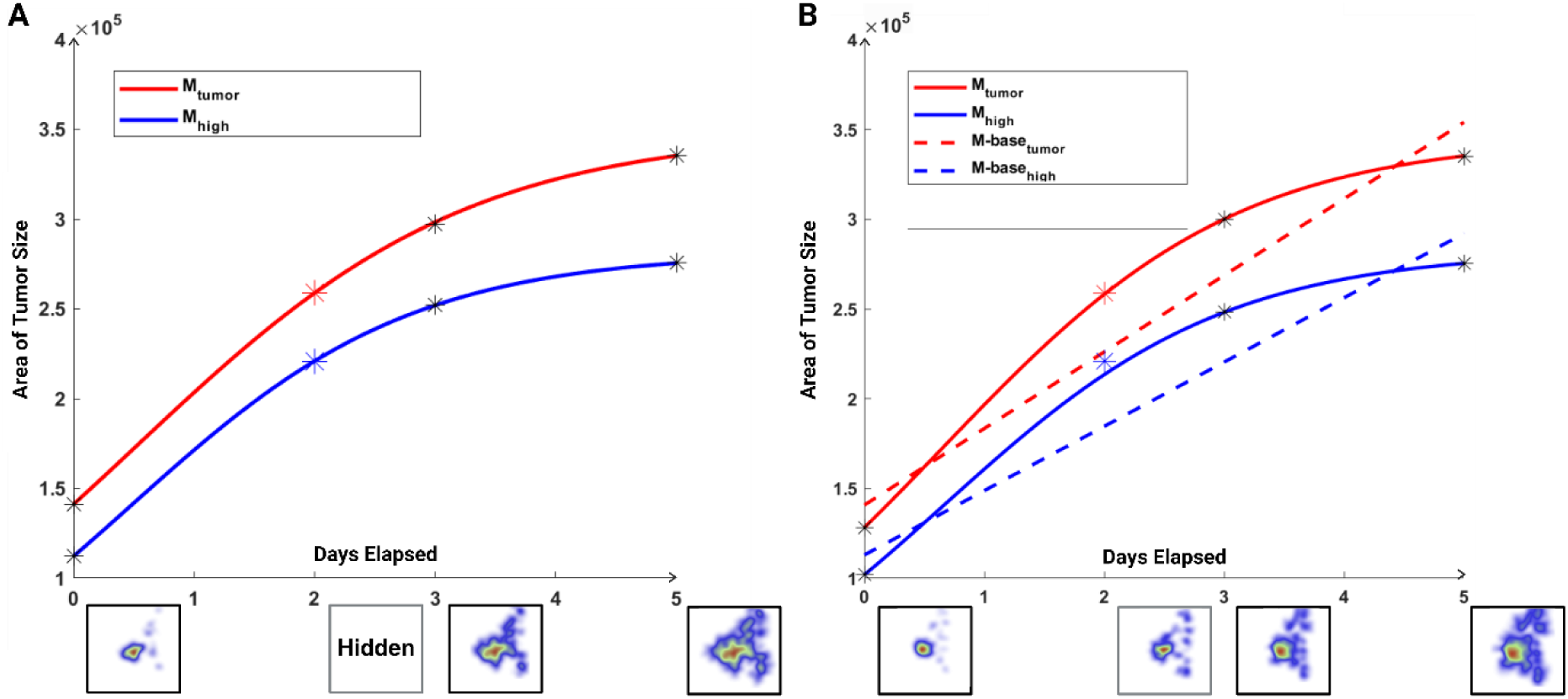
MLPA Model Testing and Validation Predictions: A) MLPA testing model simulation for tumor and high-density tumor areas over five days, with Day 2 data hidden to evaluate accuracy. Solid lines show predictions, and asterisks mark observed data from the mice dataset; and B) Validation model predictions trained on Day 0, 3, and 5 data from MLPA testing instance are compared to baseline linear regression (dotted lines) to assess error.

Then, the tumor cell density for Day 2 was analyzed using the newly calibrated parameters. These predictions were compared with the original Day 2 observations. The 95% confidence interval for the validation MLPA model’s Day 2 tumor pixel count was determined to be between 239,413 and 281,988 pixels when the actual tumor pixel count was 258,758. The 95% confidence interval for the validation MLPA model’s Day 2 high-density tumor pixel count was determined to be between 208,999 and 231,933 pixels when the actual high-density tumor pixel count was 220,884. In both cases, the 95% confidence interval of the validation MLPA model encapsulates the recorded testing MLPA model’s pixel counts, indicating that predictions for tumor cell density on Day 2 are statistically consistent with the testing data.

To further validate the model’s predictive capabilities, the analysis was extended by simulating tumor progression while hiding Day 5 data, using only the initial parameters trained from days 0, 2, and 3. In this scenario, a new validation MLPA model was trained using data from days 0, 2, and 3 from the same testing MLPA model with the same testing model parameters as before, optimizing the parameters to best fit the observed data. The training process yielded optimized parameters in the validation MLPA model of P_growth_ = 4.68 × 10^−2^ and P_metastasis_ = 2.72 × 10^−4^, as shown in Figure 4. The predicted parameters were consistent with those obtained from the validation model predicting Day 2 imaging data, showing consistency when using the same data.

**Figure 4:**
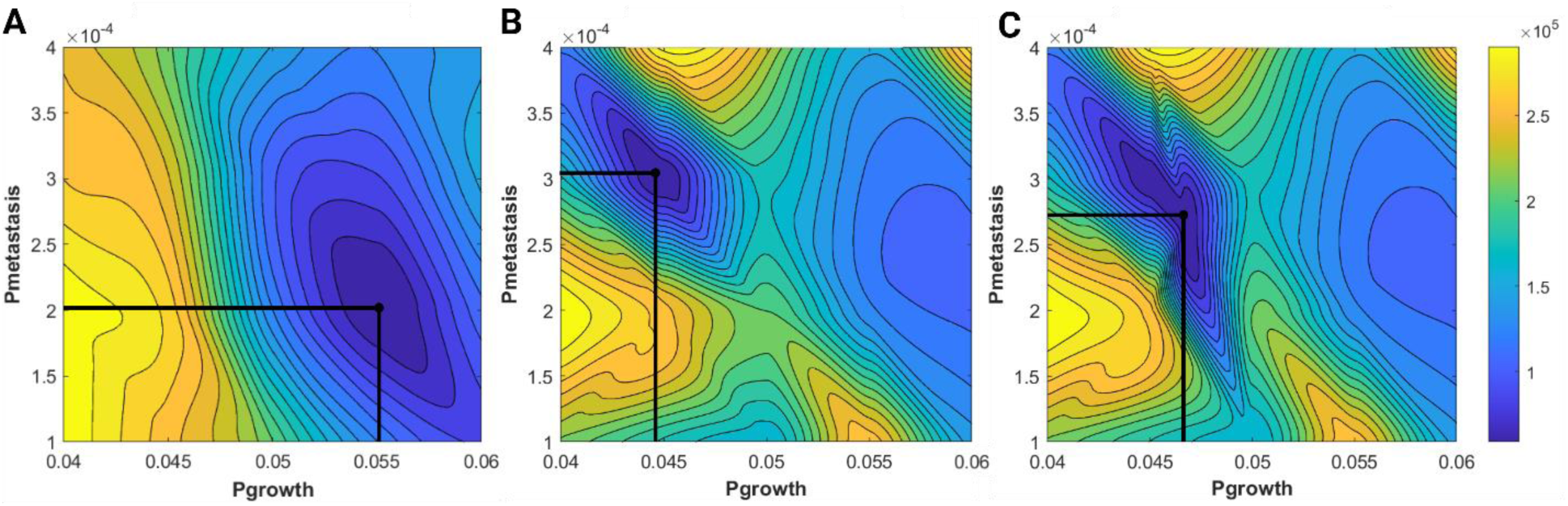
Monte Carlo simulations for MLPA parameter optimization: A) Based on the mice imaging dataset, P_growth_ = 5.5 × 10^−2^ and P_metastasis_ = 2.0 × 10^−4^; B) Based on hidden Day 2 trial, P_metastasis_ = 2.0 × 10^−4^ and P_metastasis_ = 3.06 × 10^−4^; and C) Based on the hidden Day 5 trial, P_metastasis_ = 4.68 × 10^−2^ and P_metastasis_ = 2.72 × 10^−4^.

The tumor cell density for Day 5 was predicted using the new validation MLPA model. These predictions were compared with the original Day 5 observations. The 95% confidence interval for the validation MLPA model’s Day 5 tumor pixel count was determined to be between 324,173 and 375,250 pixels when the actual tumor pixel count was 358,011. The 95% confidence interval for the validation MLPA model’s Day 5 high-density tumor pixel count was determined to be between 245,823 and 287,332 pixels when the actual high-density tumor pixel count was 257,161. In both cases, the 95% confidence interval of the validation MLPA model encapsulates the recorded testing MLPA model’s pixel counts, indicating that the model’s predictions for tumor cell density on Day 5 are statistically consistent with the original model data. This result displays the model’s reliability in predicting future time points with a predetermined set of data.

In addition, MLPA’s performance was compared with a baseline linear regression, shown in Figure 3B. These results demonstrate that MLPA’s predictions for tumor and high-density cell distributions are significantly more accurate than the baseline model, with an average tumor residual of 666 compared to the baseline’s 32,710, and an average high-density tumor residual of 7,466 compared to the baseline’s 36,269, effectively capturing the processes of tumor progression.

## Discussion

The MLPA framework demonstrates robust predictive power in simulating cancer growth and assessing treatment responses, validated with an R^2^ value of 0.93 from bioluminescent imaging data of mice with leukemia. The model achieved state-of-the-art accuracy in capturing tumor behavior under untreated, aggressive, and drug-treated conditions, with DOT1L treatment predictions consistently falling within 95% confidence intervals across multiple time points. This strong validation underscores MLPA’s capacity to simulate tumor dynamics accurately, providing insights into cancer progression and treatment efficacy.

MLPA’s approach sets it apart from traditional cancer prediction models by integrating multi-scale data, including EHRs, genomic information, and imaging datasets, allowing for the simulation of tumor behavior across multiple time points. Compared to conventional models, which often rely on static data and lack real-world calibration, MLPA leverages stochastic cellular automata and Monte Carlo simulations to explore the parameter space. This integration reduces simulation error, resulting in more reliable predictions of cancer progression. Additionally, MLPA’s capacity to simulate cancer dynamics over time offers a more comprehensive view of tumor growth, metastasis, and treatment responses.

Distinctively, MLPA incorporates gene pathway perturbations, enhancing its predictive accuracy and allowing for the exploration of genomic data in cancer simulations. This feature distinguishes it from other models, enabling the simulation of gene-targeted therapies and providing insights into how specific pathways can influence cancer progression.

By using cellular automata and Monte Carlo simulations, MLPA offers transparency into the mechanisms of tumor growth and treatment response—an advantage over black-box AI models that focus primarily on predictive metrics without explaining underlying biological factors. The framework’s transparency in parameter influence provides valuable insights for optimizing personalized treatment strategies, as shown in its virtual testbed simulations of therapeutic impacts on individual patients in Figure 5. Compared to previous AI models, MLPA not only meets predictive benchmarks but also offers clarity on how cancer progression mechanisms interact with treatment variables, supporting more informed clinical decisions.

**Figure 5:**
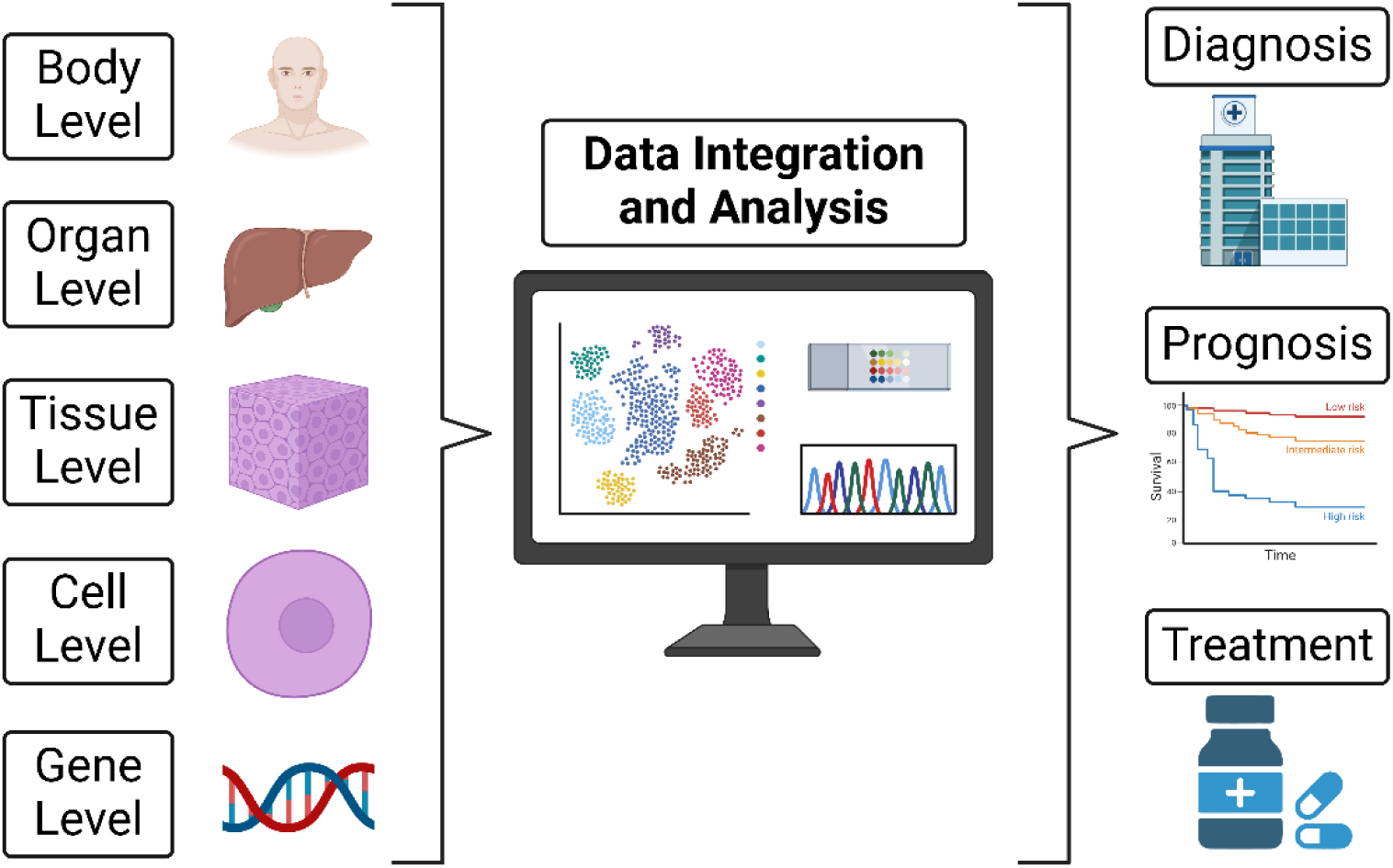
Illustration of multiscale data integration in healthcare using digital twins. Different biological levels, from the body to the gene level, are integrated into a central data analysis system. The system processes these layers of data to support diagnosis, prognosis, and therapy optimization.

In its current form, MLPA represents a significant advancement in cancer modeling, merging precision and transparency. The model’s predictive capacity and Monte Carlo optimization set a new standard in accuracy, crucial for personalized medicine. Moreover, its unique ability to simulate the effects of gene-targeted therapies introduces a new level of insight into cancer dynamics, with the potential to help clinicians test therapeutic scenarios virtually. Future development to expand the model into 3D simulations and integrate human datasets could further enhance its potential for advancing personalized cancer care and improving patient outcomes.

## Limitations

Despite the strengths of the MLPA framework, there are several limitations in the current research: 1) MLPA’s validation was based on mouse data ^22^. The absence of human-specific data, especially paired human genomic and imaging data, limits the framework’s capacity to make accurate predictions for human patients. In addition, continuous cancer progression data from human patients is unethical, as treatment is necessary. Instead, datasets with cancers that grow similarly in humans and other animals could be used. Incorporating such datasets are desirable to enhance the model’s precision and reliability in a real clinical setting. For example, transgenic mouse models such as MMTV-PyMT for breast cancer and KRAS-driven models for lung and pancreatic cancers have demonstrated tumor growth dynamics closely resembling their human counterparts, making them valuable for bridging the gap between murine and human data in predictive frameworks like MLPA ^23–25^; 2) While the MLPA framework is designed to incorporate EHR data, the use of mice imaging data, which did not include EHR data, and the incompatibility between animal EHR data and human EHR data, limited this research’s ability to validate the impact of these factors. Future research will focus on integrating and testing human EHR data to explore how these additional physiological parameters influence the model’s predictions and overall effectiveness in a clinical setting; and 3) It should be mentioned that although the current approach is implemented in 2D simulations due to being computationally efficient, the MLPA framework can be further expanded into 3D simulations. Expanding the model to 3D simulations will provide a more accurate representation of cancer growth, enabling a deeper understanding of spatial tumor interactions and treatment effects.

## Author contributions

Mr. James Gu was responsible for implementing the research work, coding the environment in MATLAB, validating results, and drafting the manuscript. Dr. Jake Y. Chen provided the scientific design, overall supervision, technical feedback, manuscript revision, and financial support to the research.

## Declaration of Generative AI and AI-assisted technologies in the writing process

Statement: During the preparation of this work the author(s) used ChatGPT in order to improve the language of the article. After using this tool/service, the author(s) reviewed and edited the content as needed and take(s) full responsibility for the content of the publication.

## STAR METHODS

### Key Resource Table

| RESOURCE | SOURCE | IDENTIFIER |
| --- | --- | --- |
| MATLAB<br>Source Code | GitHub | https://github.com/alphamind-club/MLPA/tree/main |
| Bioluminescent<br>Mice Imaging<br>Data | Dr. Christopher R. Vakoc; Wang, E., Kawaoka, S., Yu, M., Shi, J., Ni, T., Yang, W., Zhu, J., Roeder, R.G., & Vakoc, C.R. (2013). *Histone H2B ubiquitin ligase RNF20 is required for MLL-rearranged leukemia*. Proceedings of the National Academy of Sciences, 110(10), 3901-3906.<br>https://doi.org/10.1073/pnas.1301045110 | https://pubmed.ncbi.nlm.nih.gov/23412334/ |
| Gene Pathway<br>Data | UAB PAGER3 | https://discovery.informatics.uab.edu/PAGER/ |

## RESOURCE AVAILABILITY

### Lead contact

Mr. James Gu, Dr. Jake Y. Chen

### Material availability

With this article, we are providing a supplementary material file that provides information about the following:

1. Untreated mice subjects, see Figure S1 in the supplementary materials.
2. Treated mice subjects, see Figure S2 in the supplementary materials.
3. Visualization of stochastic cellular automata simulated growth, see Figure S3 in the supplementary materials.
4. Table S1 lists the gene pathways used in the MLPA framework and their relationship to the stochastic cellular automata parameters.

### Data and code availability <u>Data Availability</u>

PAGER3 is an online repository developed by the University of Alabama at Birmingham that provides access to pathways, annotated lists, and gene signatures (PAGs) related to human network biology. It offers tools for analyzing and browsing PAGs, facilitating research into gene functions and interactions. The bioluminescent imaging data was obtained from the study titled “Histone H2B ubiquitin ligase RNF20 is required for MLL-rearranged leukemia”, which investigated the role of RNF20, an enzyme that modifies histone H2B, in the development of leukemia associated with MLL gene rearrangements.

### Code Availability

The code related to MLPA is provided through the GitHub repository. The resource contains the main model script for cancer simulation, contour plots for data visualization, algorithms to find residual errors from the simulation, and dataset CSV files. The code and experiment-related materials can be obtained from the following GitHub link https://github.com/alphamind-club/MLPA/tree/main.

### Acronyms

| Acronym | Definition |
| --- | --- |
| MLPA | Multi-level Parameterized Automata |
| EHR | Electronic Health Record |
| R2 | R-Square Score |
| AI | Artificial Intelligence |
| VEGF | Vascular Endothelial Growth Factor |
| EMT | Epithelial-Mesenchymal Transition |
| CSV | Comma-separated Values |
| MLL | Mixed-lineage Leukemia |
| DOT1L | Disruptor of Telomeric Silencing 1-like |
| RGB | Red, Green, Blue |
| MAE | Mean Absolute Error |
| RMSE | Root Mean Square Error |
| PAG | Pathways, Annotated Lists, and Gene Signatures |

## METHOD DETAILS

### Novel MLPA Framework Overview

To simulate cancer progression and optimize treatment strategies, this research employs a dual-level approach, creating a personalized cancer digital twin to obtain higher prediction accuracies. At the physiological level, the MLPA framework focuses on simulating the physical aspects of cancer progression within the patient’s body using imaging and EHR data, such as sex, age, BMI, weight, blood pressure, and other factors. This includes accurately mapping the spatial location and growth dynamics of tumors, simulated using stochastic cellular automata where cells within a defined neighborhood have probabilities of becoming tumor cells based on local conditions and growth potentials. On the other hand, at the molecular level, the framework integrates genomic data to simulate critical gene pathways affecting tumor behavior, including growth, mutation, and metastasis. As shown in Figure 6A, the molecular data enables bi-directional communication with the physiological level, influencing parameters such as tumor growth probabilities and metastasis rates.

**Figure 6:**
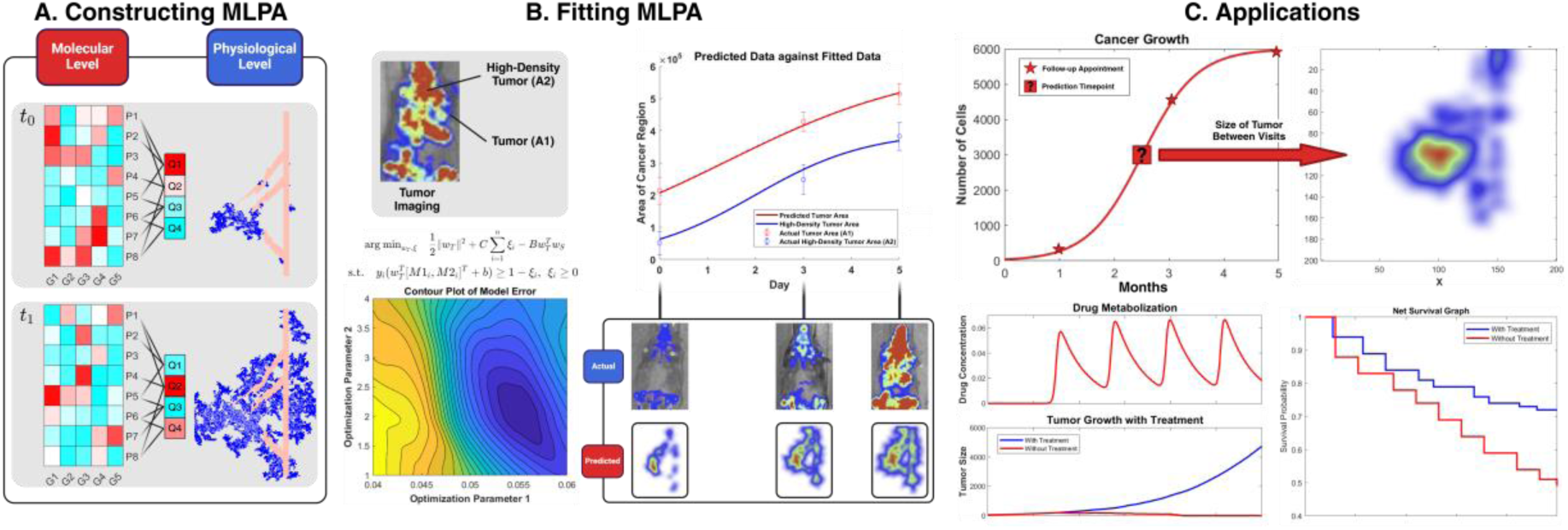
An overview of the proposed novel MLPA strategy to simulate cancer growth and metastasis: A) The framework maps gene pathways G1-G5 to model parameters Q1-Q5; B) Model fitting process calibrated against bioluminescent imaging data; and C) Applications include cancer progression, drug metabolization, and treatment effects on survival. Mice images were from a dataset ^22^.

Framework calibration is achieved by fitting the simulation outputs to real bioluminescent mice imaging data, captured over multiple time points. Bioluminescent imaging, a powerful tool in preclinical cancer studies, enables real-time, non-invasive visualization of tumor dynamics by detecting light emitted from luciferase-tagged cancer cells ^26^. This technique provides critical metrics such as cancer area and high-density tumor regions, making it invaluable for validating the MLPA framework. Monte Carlo simulations are then used to explore the parameter space, identifying optimal values that minimize error between simulated and observed data. Figure 6B shows the process that enhances the accuracy of the digital twin. Framework validation is done by comparing the continuous tumor growth predictions from the Monte Carlo simulation with the observed data from bioluminescent cancer imaging, as this imaging closely resembles the visual output of MLPA.

The digital twin strategy serves as a novel virtual platform for personalized medicine applications, facilitating the evaluation of treatment efficacies and the prediction of individualized patient responses to various therapies. The proposed MLPA approach provides valuable insight into the future progression of the disease under various treatment scenarios. Depicted in Figure 6C, the advantage of MLPA is providing a better understanding of how different therapies may impact tumor growth, patient survival, and drug metabolization over time.

### Proposed MLPA Architecture

Figure 7 shows the construction of the MLPA architecture, which involves three core components besides the base imaging component: the physiological level, the molecular level, and the AI mapping module. Each of these components plays a crucial role in simulating cancer progression and treatment responses. At the physiological level, the architecture simulates the physical aspects of cancer within the patient’s body, such as tumor growth and spread. It integrates EHR data, including variables like patient age, gender, BMI, and other relevant health metrics. The physiological component is designed to reflect how these macroscopic factors influence cancer progression, providing a broader context for the simulation. At the molecular level, MLPA focuses on simulating molecular and cellular processes, including gene pathway activity that influences tumor behavior, such as growth, mutation, and metastasis. The molecular component leverages genomic data to simulate the microscopic environment of the tumor, providing a detailed view of the internal mechanisms driving cancer progression. Due to the lack of public data correlating certain gene pathways to cancer progression, the AI mapping module serves as the bridge between the molecular level and model parameters, using ChatGPT 4o to map gene pathway relationships ^27–29^.

**Figure 7:**
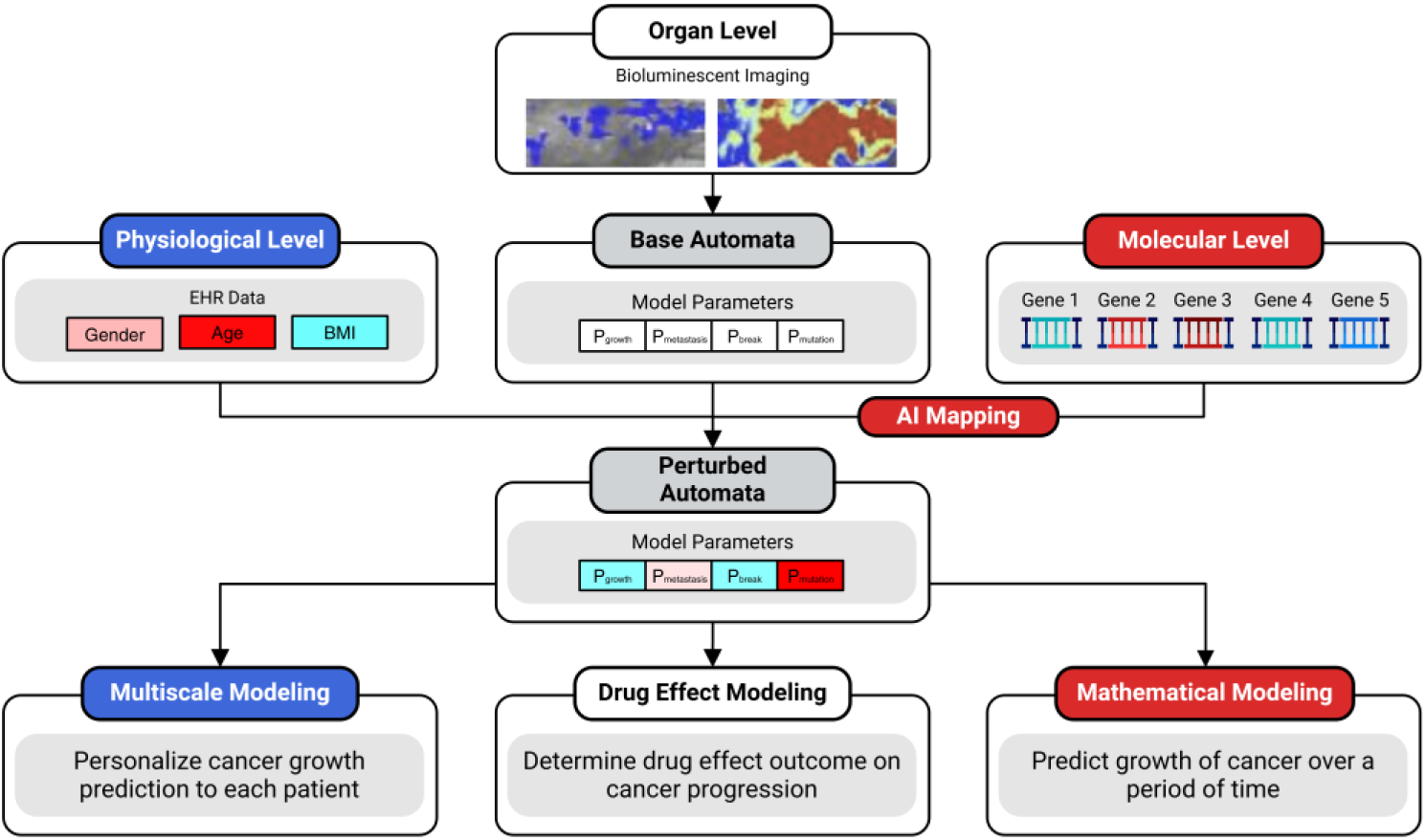
The proposed MLPA architecture integrating EHR, genomic, and imaging data mapped to model cancer progression. This multiscale approach personalizes growth predictions and simulates drug effects on tumor dynamics while considering mathematical models. Mice images were taken from a dataset ^22^.

The MLPA framework utilizes a stochastic cellular automata model to simulate cancer progression. The model, built inside the programming language MATLAB, captures the complex dynamics of tumor growth, mutation, and metastasis. The simulation environment is represented by a lattice defined as L = {(x, y) | 1 ≤ x ≤ 200, 1 ≤ y ≤ 200},initialized with a circular tumor located at the center of the lattice. Because a two-dimensional lattice is used, a type of search called Moore’s neighborhood, defined by N^M^(x_0_, y_0_) = {(x, y): |x – x_0_| ≤ r, |y – y_0_| ≤ r}, can be employed to model tumor growth dynamics. Tumor cells grow based on probabilistic rules within Moore’s neighborhood of within a distance r(u, v) ≔ max(|u_i_– v_i_|). A snapshot of the stochastic cellular automata’s simulated growth is provided in Figure S1.

Adjacent cells within the Moore’s neighborhood can become tumor cells, with a defined probability P_growth_. P_growth_ is also influenced by a growth rate map generated using a Gaussian-filtered random map with a standard deviation of 1.00. This map is normalized and polarized to create areas with varying growth potentials, effectively modeling the heterogeneous nature of the tumor microenvironment. Growth rate values range from 0.02 to 1.00, with higher growth rates assigned to regions with lower polarized map values and vice versa, as shown in Figure 8A.

**Figure 8:**
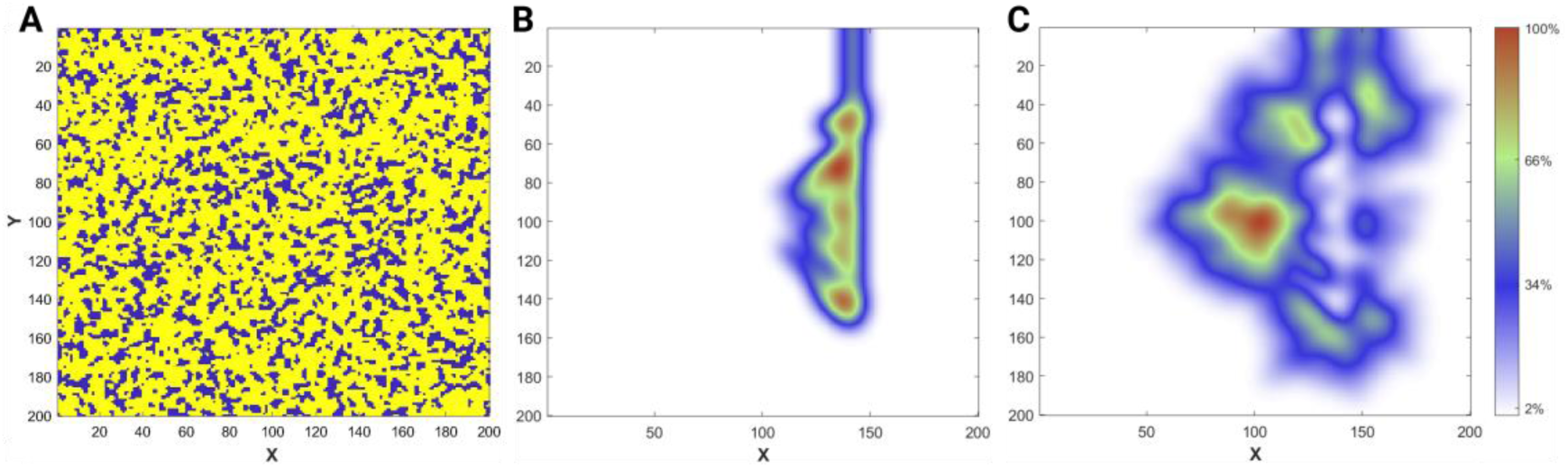
Simulation initialization and results using MLPA: A) P_growth_ is influenced by a Gaussian-filtered map which models tumor microenvironment nutrient heterogeneity; B) Blood vessel heatmap showing angiogenesis; and C) Heatmap of tumor cells expressing VEGF signaling.

Blood vessels have a chance P_break_to grow towards nearby cancer cells. Blood vessels start as rectangular regions from one lattice end. Vessel growth is directed toward Vascular Endothelial Growth Factor (VEGF) signals from tumor cells, simulating angiogenesis, highlighted in Figures 8B and 8C. The blood vessel growth follows a probabilistic rule, with potential Epithelial-Mesenchymal Transition (EMT) breaks dictated by a decreasing probability P_break_as breaks accumulate. To enhance nutrient delivery, the new EMT breaks grow toward the nearest tumor cell, guided by VEGF signals. Cells near blood vessels have their growth probability increased by 50% due to nutrient availability.

Metastasis, defined as P_metastasis_, starts when cancer cells N_cancer_surpass a defined threshold N_threshold_. The metastasis process results in new tumor sites near blood vessels, simulating cancer’s vascular spread. Additionally, tumor cells can also mutate with a probability P_mutation_, changing the cell’s P_growth_ and P_metastasis_. An example of how the parameters is used is shown in Algorithm 1 under Table 1.

**Table 1:**
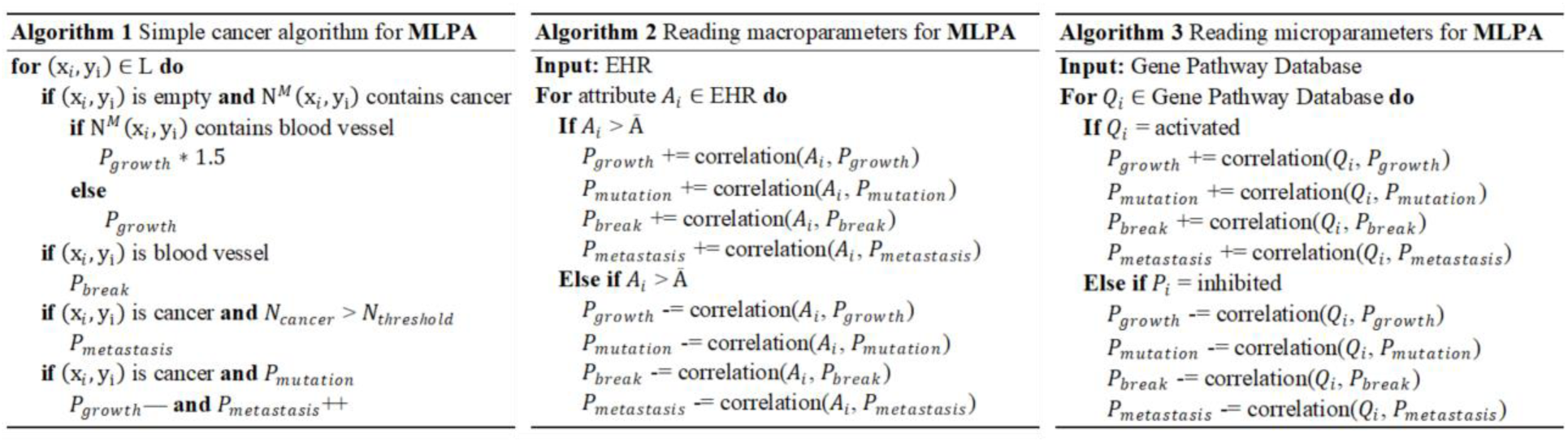
Summary of representative algorithms used in MLPA framework.

| Algorithm 1 Simple cancer algorithm for MLPA | Algorithm 2 Reading macroparameters for MLPA | Algorithm 3 Reading microparameters for MLPA |
| --- | --- | --- |
| **for** $(x_i, y_i) \in L$ **do**<br>**if** $(x_i, y_i)$ is empty **and** $N^M(x_i, y_i)$ contains cancer<br>**if** $N^M(x_i, y_i)$ contains blood vessel<br>$P_{growth} \ast 1.5$<br>**else**<br>$P_{growth}$<br>**if** $(x_i, y_i)$ is blood vessel<br>$P_{break}$<br>**if** $(x_i, y_i)$ is cancer **and** $N_{cancer} > N_{threshold}$<br>$P_{metastasis}$<br>**if** $(x_i, y_i)$ is cancer **and** $P_{mutation}$<br>$P_{growth} \text{---} \text{and } P_{metastasis} \text{++}$ | **Input:** EHR<br>**For** attribute $A_i \in \text{EHR}$ **do**<br>**If** $A_i > \bar{A}$<br>$P_{growth} \text{+= correlation}(A_i, P_{growth})$<br>$P_{mutation} \text{+= correlation}(A_i, P_{mutation})$<br>$P_{break} \text{+= correlation}(A_i, P_{break})$<br>$P_{metastasis} \text{+= correlation}(A_i, P_{metastasis})$<br>**Else if** $A_i > \bar{A}$<br>$P_{growth} \text{--= correlation}(A_i, P_{growth})$<br>$P_{mutation} \text{--= correlation}(A_i, P_{mutation})$<br>$P_{break} \text{--= correlation}(A_i, P_{break})$<br>$P_{metastasis} \text{--= correlation}(A_i, P_{metastasis})$ | **Input:** Gene Pathway Database<br>**For** $Q_i \in \text{Gene Pathway Database}$ **do**<br>**If** $Q_i$ = activated<br>$P_{growth} \text{+= correlation}(Q_i, P_{growth})$<br>$P_{mutation} \text{+= correlation}(Q_i, P_{mutation})$<br>$P_{break} \text{+= correlation}(Q_i, P_{break})$<br>$P_{metastasis} \text{+= correlation}(Q_i, P_{metastasis})$<br>**Else if** $P_i$ = inhibited<br>$P_{growth} \text{--= correlation}(Q_i, P_{growth})$<br>$P_{mutation} \text{--= correlation}(Q_i, P_{mutation})$<br>$P_{break} \text{--= correlation}(Q_i, P_{break})$<br>$P_{metastasis} \text{--= correlation}(Q_i, P_{metastasis})$ |

To personalize these model parameters to each individual patient, the incorporation of digital twin technology integrates macroscopic patient EHR and microscopic patient genomic data to create a highly individualized disease progression system. The macroscopic aspect of MLPA utilizes simulated EHR data. Blood pressure can be continuously monitored to assess the patient’s cardiovascular health, influencing tumor growth, and treatment response ^30^. Patient age and gender specific differences are factored to account for their roles in cancer progression and treatment effectiveness ^31–32^. Other EHR parameters, such as BMI and medical history, are included to comprehensively view the patient’s health status. The macroscopic data can directly perturb the stochastic cellular automata parameters such as P_growth_, P_break_, P_metastasis_, and P_mutation_, as shown in Algorithm 2 on Table 1.

On the molecular level, the genomic pathway data then perturbs the model parameters to tailor the framework to individual patients, effectively customizing the simulation based on each patient’s specific genomic profile ^33–35^. The perturbation process is guided by the PAGER3 database, which includes pathways scientifically validated to affect metastasis, growth, mutation, and angiogenesis ^36–39^. The extracted pathway data was placed into a CSV file that lists gene pathway names and their interactions to the model parameters. Pathway expression is controlled via a graphical user interface, allowing adjustments based on activation, absence, or inhibition of specific gene pathways. These adjustments directly influence the stochastic cellular automata parameters, as described by their correlation in the CSV file. For instance, an expressed gene pathway associated with increased P_growth_will adjust the growth rate parameter accordingly, while pathways involved in inhibiting P_growth_will decrease it, as shown in Algorithm 3 in Table 1.

The AI mapping module was crucial in identifying relationships between specific gene pathways and the four key model parameters, as there are no datasets available to describe the relation between specific pathways to the model parameters that we defined in MLPA. To accomplish this, ChatGPT 4o was utilized, leveraging its capabilities in natural language processing and data interpretation to evaluate potential interactions between gene pathways and model parameters while minimizing risks of AI hallucination. ChatGPT 4o was prompted with clear, pathway-specific details to determine whether each pathway had a positive, negative, or neutral effect on the parameters. Recent studies into ChatGPT’s application in biomedical contexts further underscore its effectiveness in identifying gaps within gene pathway annotations and enriching pathway data, enhancing the accuracy of this mapping process ^27–29^. Given the absence of genomic data in the dataset, a standardized approach was applied by assigning a 10% change to the parameters for any pathway relationship identified by ChatGPT, ensuring MLPA could reflect gene pathway influences on tumor behavior even without direct genomic data. A list of gene pathways implemented is available in Table S1.

### Data Acquisition and Preprocessing

The imaging dataset was sourced from the study ^22^. The dataset focuses on Mixed-lineage leukemia (MLL) fusions, potent oncogenes that initiate aggressive forms of acute leukemia. MLL-fusion proteins, as aberrant transcriptional regulators, alter gene expression in hematopoietic cells by interacting with the histone H3 lysine 79 methyltransferase DOT1L. Notably, treating leukemia using a DOT1L treatment decreased the spread of the cancer.

As described in Table 2, the dataset includes bioluminescent imaging data from 16 mice, each recorded at three time points: days 0, 3, and 5. Among these, four mice were treated with a small-molecule inhibitor of DOT1L. A 50/50 training/testing split was employed, using data from 8 untreated mice for framework training and data from the remaining 8 mice for testing (4 treated and 4 untreated). In total, 48 images were processed, each tagged with metadata corresponding to the original mouse subject. The images were standardized in size and underwent segmentation based on RGB values to distinguish different regions of the bioluminescent imaging heatmap. For each image, two critical variables were extracted: tumor pixel count (all color values) and high-density tumor pixel count (red color values only). Examples of the data are given in Figure S2 and S3.

**Table 2:** Summary of untreated and treated mice imaging data split into training and testing category.

| Mice Group | Number of Mice | Time Points | Imaging Type | Treatment Type |
| --- | --- | --- | --- | --- |
| Untreated (Training) | 8 | Days 0, 3, 5 | Bioluminescent | Untreated |
| Untreated (Testing) | 4 | Days 0, 3, 5 | Bioluminescent | Untreated |
| Treated (Testing) | 4 | Days 0, 3, 5 | Bioluminescent | DOT1L Inhibitor |

### Training Using Monte Carlo Simulations

From the bioluminescent imaging dataset, third-degree polynomials were fitted to the tumor pixel count and high-density tumor pixel count. Harshe et al. states that three tumor volume measurements are sufficient to estimate model parameters when noise is absent ^40^. As such, Day 0, Day 3, and Day 5 imaging data were used to fit the polynomial curves. These curves provided a mathematical comparison of the observed growth patterns in the bioluminescent images. Monte Carlo simulations, which are computational techniques that use repeated random sampling to estimate numerical results, were utilized ^41^. In the context of MLPA, a Monte Carlo simulation helps explore the range of possible tumor growth scenarios by varying P_growth_ and P_metastasis_ to find which parameter pair most closely represents the cancer growth in the mice dataset. A systematic approach using Monte Carlo simulations identified these ranges as optimal for minimizing residual errors in tumor pixel predictions. These ranges consistently captured the tumor growth and metastasis dynamics across control, aggressive, and treatment scenarios. Random values for P_growth_ and P_metastasis_ were chosen from their respective distributions and fed into the stochastic automata model. P_break_ and P_mutation_ were held constant with P_break_ = 1.0 × 10^−3^ and P_mutation_ = 1.0 × 10^−2^, as the bioluminescent imaging dataset did not include blood vessel mapping or genomic data to indicate mutations. The pathways from the CSV file, such as mRNA splicing, Processing of Capped Intron Containing Pre mRNA, mRNA Processing, Nonsense Mediated Decay, Chaperon Mediated Autophagy, Regulation of TP53 Expression and Degradation, and others, were assigned their typical expression in cancerous conditions, with each pathway being assigned a value of 1, -1, or 0. In this context, a value of 1 indicates that the pathway is activated, meaning it is upregulated or more active in the cancerous state. A value of -1 signifies that the pathway is inhibited, reflecting downregulation or reduced activity in the cancerous state. Lastly, a value of 0 indicates that the pathway is not expressed or has no relationship to the parameters in the stochastic cellular automata model. The AI mapping module remained active throughout this process, ensuring that these values were accurately perturbing the model parameters.

The automata’s output was segmented to match the mice images, providing pixel counts for tumor and high-density areas. Simulated pixel counts (derived from MLPA) were compared to observed pixel counts (derived from mice dataset), calculating residuals for each simulation, with errors quantified using Mean Absolute Error and Root Mean Squared Error. Error values were mapped to their respective P_growth_ and P_metastasis_ values, with the optimal values determined to be P_growth_ = 5.5 × 10^−2^ and P_metastasis_ = 2.0 × 10^−4^ by identifying the region with the lowest error values, as shown in Figure 9. These optimal values became MLPA’s default parameters, ensuring accurate tumor growth modeled after the mice dataset. By fitting the parameters of MLPA to the observed data, the system is optimized to accurately reflect the tumor growth dynamics observed in the imaging dataset.

**Figure 9:**
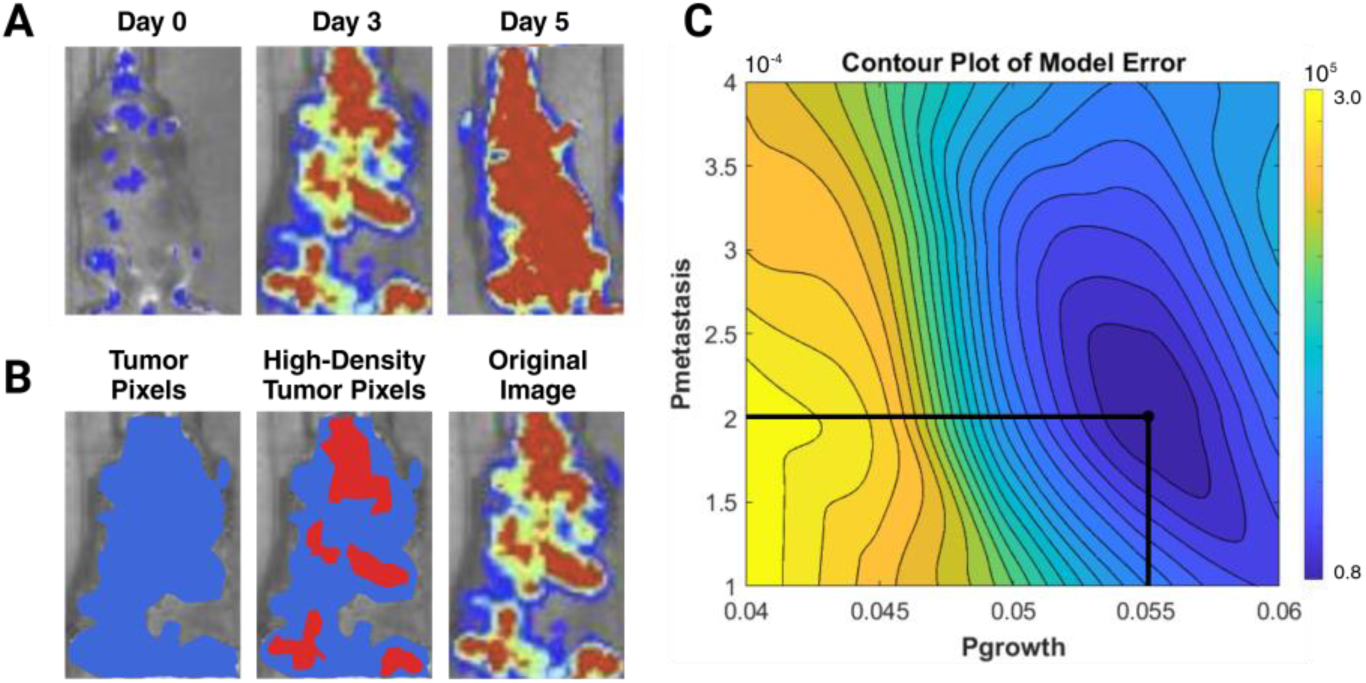
Fitting MLPA to bioluminescent imaging data and optimizing model parameters. A) Time series data consisting of day 0, day 3, and day 5; B) Segmentations of original image into tumor pixels and high-density tumor pixel segmentations; and C) Contour plot of prediction error used to fine-tune model parameters, with P_growth_ = 5.5 × 10^−2^ and P_metastasis_ = 2.0 × 10^−4^ set as default. Mice images were obtained from a dataset ^22^.

