## Supplemental File for "Creation of MLPA: A Multi-level Digital Twin Framework for Personalized Cancer Simulation and Treatment Optimization"

Supplemental Items

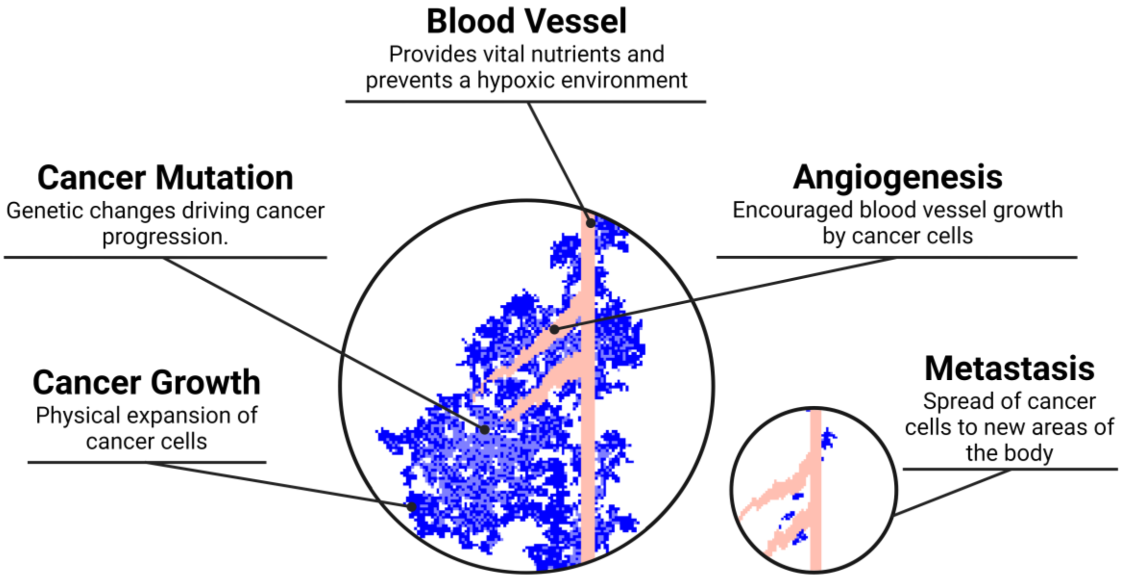


Figure S1: Illustration of tumor progression within the MLPA simulation environment, highlighting key processes and spatial interactions. The pink structures represent angiogenesis pathways, modeling the formation of new blood vessels to supply nutrients to tumor cells. The blue regions denote tumor cells, which can mutate, turning light blue and adopting different growth parameters.

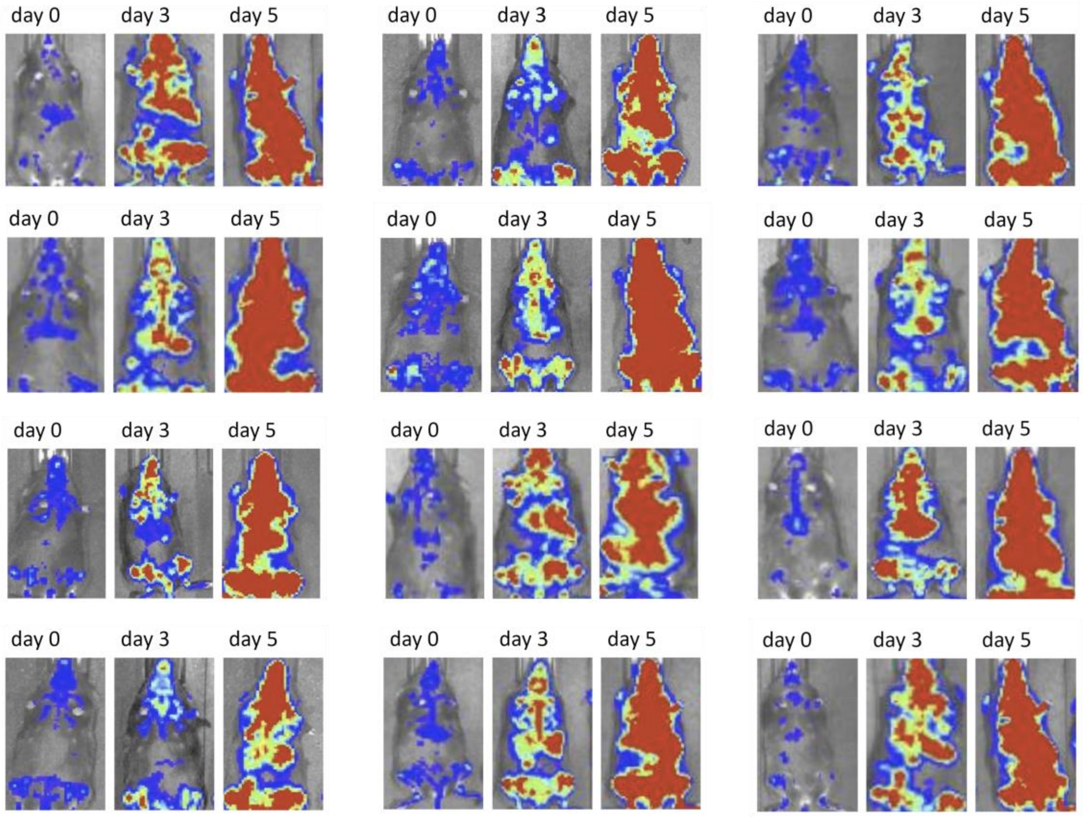


Figure S2: Representative bioluminescent imaging data showing tumor progression in 12 mice subjects at three time points: day 0, day 3, and day 5. Each panel illustrates the tumor region in a grayscale image overlaid with bioluminescent signal intensity, where blue represents low intensity, yellow represents medium intensity, and red represents high intensity. The progression of tumor growth and increasing bioluminescent signal intensity from day 0 to day 5 indicates the dynamic expansion of tumor regions. Mice images were from dataset [S1].

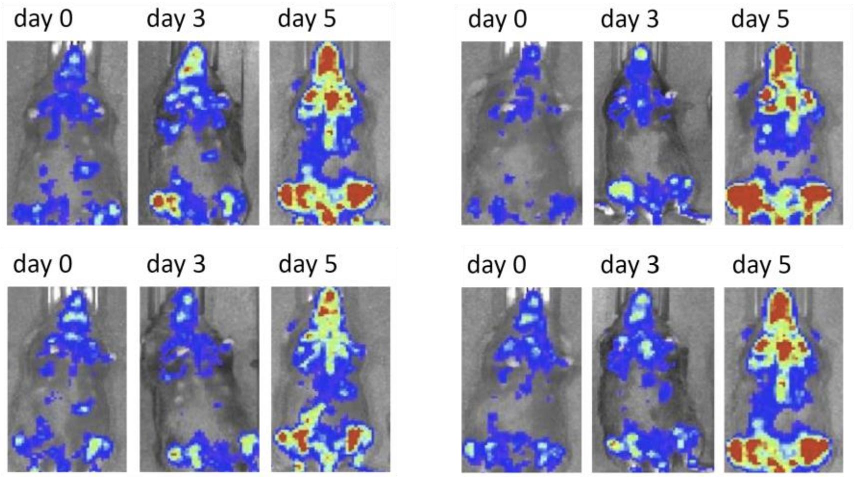


Figure S3: Representative bioluminescent imaging data showing tumor progression in 4 mice models subjected to treatment at three time points: day 0, day 3, and day 5. Each panel displays grayscale images overlaid with bioluminescent signal intensity, where blue indicates low signal, yellow indicates moderate signal, and red indicates high signal. The treated group exhibits reduced tumor growth and signal intensity compared to untreated models, demonstrating the efficacy of the therapeutic intervention. Mice images were from dataset [S1].

| Pathway Name | Pbreak | Pgrowth | Pmutation | Pmetastasis |
|---|---|---|---|---|
| Genes Involved in mRNA Splicing | + | + | + | ? |
| Genes Involved in Processing of Capped Intron Containing Pre mRNA | + | + | ? | + |
| Genes Involved in mRNA Processing | + | + | N/A | + |
| Formation of an Intermediate Spliceosomal C (Bact) Complex | + | + | ? | + |
| Formation of Exon Junction Complex | ? | ? | ? | ? |
| Cleavage at the 3' Splice Site and Exon Ligation | + | + | N/A | + |
| Formation of the Spliceosomal A Complex | ? | ? | ? | ? |
| Formation of the Spliceosomal B Complex | ? | ? | ? | ? |
| Formation of the Active Spliceosomal C (B*) Complex | ? | ? | ? | ? |
| Lariat Formation and 5' Splice Site Cleavage | + | + | N/A | + |
| Nonsense Mediated Decay | - | - | N/A | - |
| Chaperon Mediated Autophagy | - | - | N/A | - |
| Regulation of TP53 Expression and Degradation | - | - | - | - |
| TP53 activity through methylation | - | - | - | - |
| Pexophagy | ? | - | ? | ? |
| Aggrephagy | - | - | ? | - |

Table S1: Gene pathways identified using the PAGER 3.0 database are mapped to stochastic cellular automata parameters, including tumor growth, mutation, angiogenesis, and metastasis. Rows represent gene pathways, and columns denote the corresponding model parameters. Gene pathways were found through the Pager 3.0 database [S2-S5]
